# Global MEG Resting State Functional Connectivity in Children with Autism and Sensory Processing Dysfunction

**DOI:** 10.1101/2024.01.26.577499

**Authors:** Carly Demopoulos, Xuan Jesson, Molly Rae Gerdes, Barbora G. Jurigova, Leighton B. Hinkley, Kamalini G. Ranasinghe, Shivani Desai, Susanne Honma, Danielle Mizuiri, Anne Findlay, Srikantan S. Nagarajan, Elysa J. Marco

## Abstract

Sensory processing dysfunction not only affects most individuals with autism spectrum disorder (ASD), but at least 5% of children without ASD also experience dysfunctional sensory processing. Our understanding of the relationship between sensory dysfunction and resting state brain activity is still emerging. This study compared long-range resting state functional connectivity of neural oscillatory behavior in children aged 8-12 years with autism spectrum disorder (ASD; N=18), those with sensory processing dysfunction (SPD; N=18) who do not meet ASD criteria, and typically developing control participants (TDC; N=24) using magnetoencephalography (MEG). Functional connectivity analyses were performed in the alpha and beta frequency bands, which are known to be implicated in sensory information processing. Group differences in functional connectivity and associations between sensory abilities and functional connectivity were examined. Distinct patterns of functional connectivity differences between ASD and SPD groups were found only in the beta band, but not in the alpha band. In both alpha and beta bands, ASD and SPD cohorts differed from the TDC cohort. Somatosensory cortical beta-band functional connectivity was associated with tactile processing abilities, while higher-order auditory cortical alpha-band functional connectivity was associated with auditory processing abilities. These findings demonstrate distinct long-range neural synchrony alterations in SPD and ASD that are associated with sensory processing abilities. Neural synchrony measures could serve as potential sensitive biomarkers for ASD and SPD.

## Introduction

Sensory dysfunction is estimated to impact at least 70% of individuals with Autism Spectrum Disorders (ASD; Adamson, Hare, & Graham, 2006; Al-Heizan, AlAbdulwahab, Kachanathu, & Natho, 2015; Greenspan & Wieder, 1997; Mayes & Calhoun, 1999; Tomcheck & Dunn, 2007), and with its recognition as a core symptom in DSM-5 (American Psychiatric Association 2013), there is a rapidly growing body of research focused on understanding the causes and impact of sensory dysfunction in ASD. This line of research can be advanced not only by studying sensory dysfunction in individuals with ASD and other clinical populations, but also through examination of the estimated >5% of non-autistic individuals who experience clinically significant sensory processing dysfunction (SPD) (Ahn et al. 2004). Yet, despite the impairment in adaptive functioning associated with SPD, the absence of a recognized categorical diagnosis limits access to resources for research and treatment in affected individuals.

Nevertheless, biological differences, such as white matter abnormalities (Chang et al. 2014; Owen et al. 2013) and cortical response latencies (Demopoulos et al. 2017), have been identified in children with SPD and these measurable structural and physiologic differences have been associated with sensory processing behaviors (Chang et al., 2016). While some features of sensory dysfunction may be shared among children with SPD and those with ASD, such as tactile processing deficits (Demopoulos, Brandes-Aitken, et al. 2015), some domains of sensory dysfunction may identify important distinctions between these populations. For example, auditory processing abnormalities have been identified as distinguishing ASD from SPD groups in both behavioral tasks and neural response latencies (Demopoulos et al. 2017; Demopoulos, Brandes-Aitken, et al. 2015). Understanding these similarities and differences in sensory processing dysfunction among children with and without ASD can not only help delineate the sensory dysfunction that is specific to ASD, but it can also heighten our understanding of sensory information processing more broadly and guide treatment strategies.

Because differences in resting state oscillatory activity can be indicative of functional pathology (Papanicolaou 2009), there has been extensive research examining differences in resting state brain activity in individuals with and without ASD diagnoses. While previous sensory processing research has focused on differences in performance-based measures of, and neural responses to, sensory processing (Chang et al., 2014; Demopoulos et al., 2015, 2017), our understanding of the relationship between sensory dysfunction and resting state brain activity is still emerging. This study will be the first to use using silently acquired recording via magnetoencephalography (MEG) to examine whole brain functional connectivity during rest in participants with ASD, SPD, and typically developing children (TDC). The goal of this study is to identify relevant differences in whole brain functional connectivity that may be associated with sensory dysfunction. Concurrent examination of these three groups offers two key benefits. First, it will add to the emerging literature identifying the shared and distinct patterns of neural activity in children with ASD and SPD. Second, it will allow us to examine differences in functional connectivity and behavioral measures of sensory discrimination in affected children. Prior research has suggested that auditory and tactile processing are particularly impacted in children with ASD (Fernandez-Andres et al. 2015), and that auditory processing has been associated with the communication impairments that are core to ASD (Demopoulos et al. 2017; Demopoulos, Brandes-Aitken, et al. 2015; Demopoulos, Hopkins, et al. 2015; Edgar et al. 2013, 2014; Lerner, McPartland, and Morris 2013; Oram-Cardy et al. 2005; Oram Cardy et al. 2008; Roberts et al. 2011, 2012, 2019, 2008, 2010). As such, we also examine associations between functional connectivity and performance-based measures of auditory and tactile processing and verbal abilities.

Our functional connectivity analyses were performed in the alpha and beta frequency bands, which are known to be implicated in sensory information processing. Specifically, these frequency bands have been associated with sensory gating (Buchholz, Jensen, and Medendorp 2014) and direction of sensory attention in the auditory and visual cortex for alpha (Foxe and Snyder 2011) and in the somatosensory cortex for beta (Bauer, Kennett, and Driver 2012; van Ede, Jensen, and Maris 2010). Further, the role of alpha activity in states of psychological distress has been widely studied (Adolph and Margraf 2017; Boutcher and Landers 1998; Demerdzieva and Pop-Jordanova 2015; Fingelkurts et al. 2007; Knyazev, Savostyanov, and Levin 2006; Mennella, Patron, and Palomba 2017; Smith, Zambrano-Vazquez, and Allen 2016), and may be relevant to differences in psychological response to sensory input in our clinical groups.

Prior research has demonstrated that both children with SPD and ASD were impaired on behavioral and neural measures of tactile processing, but only the ASD group demonstrated auditory dysfunction (Demopoulos et al. 2017; Demopoulos, Brandes-Aitken, et al. 2015). This work is consistent with structural findings that children with ASD and SPD demonstrate decreased connectivity in parieto-occipital tracts, but connectivity in temporal tracts was only reduced in the ASD group (Chang et al., 2014). Thus, given these shared and divergent sensory findings between children with ASD and SPD, and given that alpha and beta connectivity has been associated with sensory gating and sensory attention in these frequency bands (Buchholz et al. 2014; Foxe and Snyder 2011), we hypothesize that similar shared and divergent MEG-derived findings of resting state functional connectivity in the alpha and beta ranges will be identified between children with ASD, SPD, and TDC participants. In addition, based on work implicating alpha oscillations in the direction of auditory attention (Bauer et al. 2012) and evidence of somatosensory cortex beta band modulation in advance of tactile stimuli (van Ede et al. 2010), we also hypothesize that alpha connectivity will be associated with auditory processing and beta connectivity will be associated with tactile processing. To test these hypotheses, these frequency bands were subjected to source space reconstruction for analysis of differences in long-range neural synchrony and associations with sensory processing abilities.

## Methods

### Participants

Participants were 60 boys aged 8-12 years (ASD N=18; SPD N=18; typically developing controls (TDC) N=24) who were recruited from the UCSF Sensory Neurodevelopmental and Autism Program (SNAP) participant registry and website, UCSF SNAP clinic, and local online parent groups. Experimental protocols were approved by the UCSF IRB and carried out in accordance with those approved procedures. Participants provided their written assent and written informed consent was obtained from parents or legal guardians prior to enrollment.

Consent and assent procedures were witnessed by a member of the study team. Participants were recruited between 5/22/2003 and 10/26/2015. All participants who were taking medication were on a stable dose for at least six weeks prior to testing as reported in our previously published studies that recruited from this pool of participants (Demopoulos et al. 2017; Demopoulos, Brandes-Aitken, et al. 2015). Specifically, in the TDC group one participant regularly used an antihistamine and a leukotriene inhibitor for seasonal allergies as well as melatonin for sleep.

Another TDC participant regularly used steroid medications paired with a bronchodilator as needed for asthma and allergies and omeprazole for reflux. A third TDC participant regularly used methylphenidate for attention. In the SPD group, one participant was prescribed lisdexamfetamine, sertraline, and valproic acid for inattention and challenging behavior, and four others were taking stimulants (amphetamine/dextroamphetamine and methylphenidate) for inattention. One additional SPD participant was taking nonstimulant medication (atomoxetine) for inattention and montelukast for allergies, and another was taking steroid medication for asthma. In the ASD group, one participant was taking a chelation agent (DMSA), another participant was taking escitalopram for anxiety, and a third was taking guanfacine and methylphenidate for calming and inattention.

Inclusion/exclusion criteria and diagnostic classification followed the criteria utilized in previous studies (Demopoulos et al. 2017; Demopoulos, Brandes-Aitken, et al. 2015). Specifically, exclusion criteria included (1) bipolar disorder, psychotic disorder, or other neurological disorder or injury, and (2) a score of 70 or below on the Wechsler Intelligence Scale for Children-Fourth Edition (WISC-IV; Wechsler, 2003) Perceptual Reasoning Index (PRI). The PRI rather than the Full Scale Intelligence Quotient (FSIQ) was utilized for exclusion criteria because verbal abilities (represented in the Verbal Comprehension Index and incorporated into the FSIQ) were examined as an outcome measure in this study. Specifically, those with prior clinical diagnosis of ASD and those scoring ≥15 on the Social Communication Questionnaire (SCQ; Rutter, Bailey, & Lord, 2003), regardless of previous diagnostic status, were evaluated with the Autism Diagnostic Inventory-Revised (ADI-R; Lord, Rutter, & Le Couteur, 1994) and the Autism Diagnostic Observation Schedule (ADOS; Lord et al., 1989). Diagnostic cutoffs on both of these measures were met for participants in the ASD group, who also met DSM-IV-TR criteria for Autistic Disorder, confirmed by a pediatric neurologist (EJM). SPD participants were previously diagnosed with SPD by a community occupational therapist. Inclusion criteria for this group were included (1) SCQ score <15 and (2) a score in the “Definite Difference” range in one or more of the auditory, visual, oral/olfactory, tactile, vestibular, or multisensory processing domains of the Sensory Profile (Dunn 1999). All SCQ and Sensory Profile scores for the TDC group were not in clinical ranges. Demographic characteristics of the study sample are presented in Table 1.

**Table 1.**
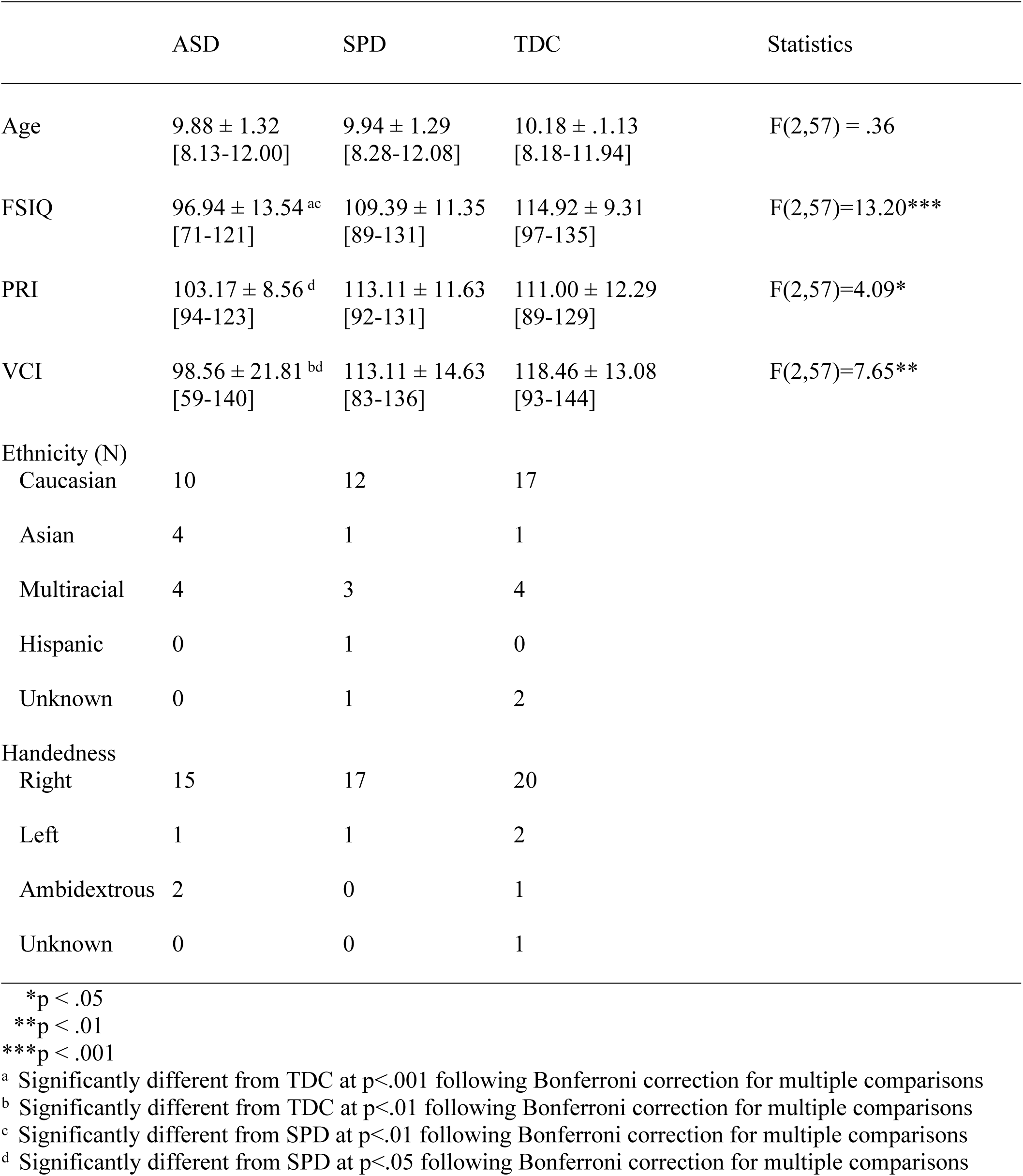
Group Characteristics (M ± SD [range])

### Measures

#### Tactile Processing

Tactile processing measures were assessed according to previously published procedures (Demopoulos et al. 2017; Demopoulos, Brandes-Aitken, et al. 2015).

Tactile form discrimination was assessed using the Van Boven Domes task (Van Boven & Johnson, 1994) and quantified by the lowest grating size of passed trials. Tactile proprioception was measured according to the total score of the right and left hand scores of the graphesthesia subtest of the Sensory Integration Praxis Tests (Ayres 1989).

#### Auditory Processing

Auditory processing also was assessed according to previously published procedures (Demopoulos, Brandes-Aitken, et al. 2015; Demopoulos et al., 2017) via the Acoustic (AI) and Acoustic-Linguistic Index (ALI) of the Differential Screening Test for Processing (DSTP; Richard & Ferre, 2006). The AI is derived from performance on measures of dichotic listening, temporal sequencing, and auditory filtering skills. The ALI assesses auditory processing skills associated with language via tasks focused on phonic and phonemic manipulation.

#### Verbal Abilities

Because auditory processing dysfunction has been repeatedly associated with weaker verbal abilities in children with ASD (Demopoulos et al. 2017; Edgar et al. 2013; Oram-Cardy et al. 2005; Roberts et al. 2011; Russo et al. 2009; Schmidt et al. 2009), we also assessed for associations between functional connectivity and verbal abilities in the ASD group using established protocols for our assessment of verbal abilities (Demopoulos et al. 2017; Demopoulos, Brandes-Aitken, et al. 2015). The Linguistic Index (LI) of the DSTP was used to evaluate semantic and pragmatic aspects of language. The VCI of the WISC-IV (Wechsler, 2003) was used to index verbal intellectual abilities.

#### Magnetic Resonance Image (MRI) Acquisition and Processing

Structural MRIs were acquired for co-registration with MEG functional data on a 3T Siemens MRI scanner at the UCSF Neuroscience Imaging Center. T1-weighted images were spatially normalized to the standard Montreal Neurological Institute template brain using 5mm voxels in SPM8 (http://www.fil.ion.ucl.ac.uk/spm/software/spm8/). Normalization results were manually verified in all participants.

#### Magnetoencephalographic Image Acquisition and Processing

Methods for acquisition and processing of MEG data follow protocols similar to those used in prior research employing these imaginary coherence metrics (Demopoulos et al. 2020; Ranasinghe et al. 2017). Specifically, MEG data were acquired at a 1200 Hz sampling rate using a 275-channel CTF System whole-head biomagnetometer (MEG International Services Ltd., Coqiotlam, BC, Canada). Fiducial coils were placed at the nasion and bilateral peri-auricular points to localize the head to the sensor array. These localizations were utilized for coregistration to the T1- weighted MRI and generation of a head shape. Four minutes of continuous recording was collected from each subject while awake with eyes closed in a supine position. While keeping eyes closed can increase alpha in resting state activity, it also serves to control visual stimulation and because this procedure was implemented for all participants, this would not confound group contrasts. As such, we elected to use an eyes closed approach, as has been used in many previous studies of resting state activity in children with ASD (Berman et al. 2015; Brodski-Guerniero et al. 2018; Cornew et al. 2012; Edgar et al. 2019; Edgar, Heiken, et al. 2015; Green et al. 2020, 2022; Port et al. 2019). Based on previous studies demonstrating reliable results from 60 second segments of MEG resting state data (Guggisberg et al. 2008; Hinkley 2010; Hinkley et al. 2011), we selected a 60-second artifact-free epoch. Artifact rejection criteria were signal amplitude >10pT or visual evidence of movement or muscle contractions.

A whole brain lead field was computed according to a spatially normalized MRI with a 10mm voxel size. The Neurodynamic Utility Tool for MEG (NUTMEG; http://nutmeg.berkeley.edu; Dalal et al., 2011) was used for source-space reconstruction and functional connectivity analyses. Source-space was reconstructed from filtered sensor (fourth-order Butterworth filter of 1–20 Hz). A linear combination of spatial weighting and sensor data matrices were used to estimate each voxel’s amplitudes (Hinkley et al., 2011).

Following source space reconstruction, functional connectivity analysis was performed by computing imaginary coherence. The imaginary coherence approach excludes zero- or π- phase-lag-connectivity to eliminate neural synchrony attributable to volume spread (Nolte et al. 2004). This approach has been documented as a reliable method for estimating long-range neural synchrony (Engel et al. 2013; Guggisberg et al. 2008; Martino et al. 2011; Nolte et al. 2004), and has been shown to reduce overestimation (Guggisberg et al. 2008; Martino et al. 2011; Nolte et al. 2004). Imaginary coherence values were transformed to Fisher’s Z prior to calculating associations between each voxel and all other voxels. These associations were averaged within each voxel to derive voxel wise global connectivity values for group contrasts in the alpha and beta frequency bands. Correlations also were performed between behavioral measures and global connectivity values at each voxel for the combined group study sample. All voxel-wise results with uncorrected p < 0.05 were further subjected to a 5% False Discovery Rate multiple comparisons correction (Benjamini and Hochberg 1995) and a 5-voxel cluster correction.

#### Missing Data

Data from the sensory battery tasks are missing for some participants because these tasks were added to the protocol after these participants were enrolled. Thus, these data can be considered missing at random. DSTP data was available for 17 ASD participants, 17 TDC participants, and 11 SPD participants. Van Boven Domes were administered to 16 participants in the ASD group, 16 in the TDC group, and 11 in the SPD group. Graphesthesia was administered to 17 ASD participants, 15 TDC participants, and 11 SPD participants.

## Results

### Group Contrasts in Alpha Connectivity

Group contrasts in alpha coherence indicated that, relative to TDC participants, the ASD group showed reduced connectivity in the left fusiform and inferior occipital gyri and cerebellum and increased connectivity in the right pre- and postcentral gyri. No significant differences were identified between the ASD and SPD groups; however, the SPD group showed increased connectivity compared to TDC participants in the left middle and superior frontal gyri and in the right inferior frontal gyrus, precuneus, and inferior and superior parietal lobules. Alpha contrast results are presented in Figure 1 and summarized in Table 2.

**Figure 1.**
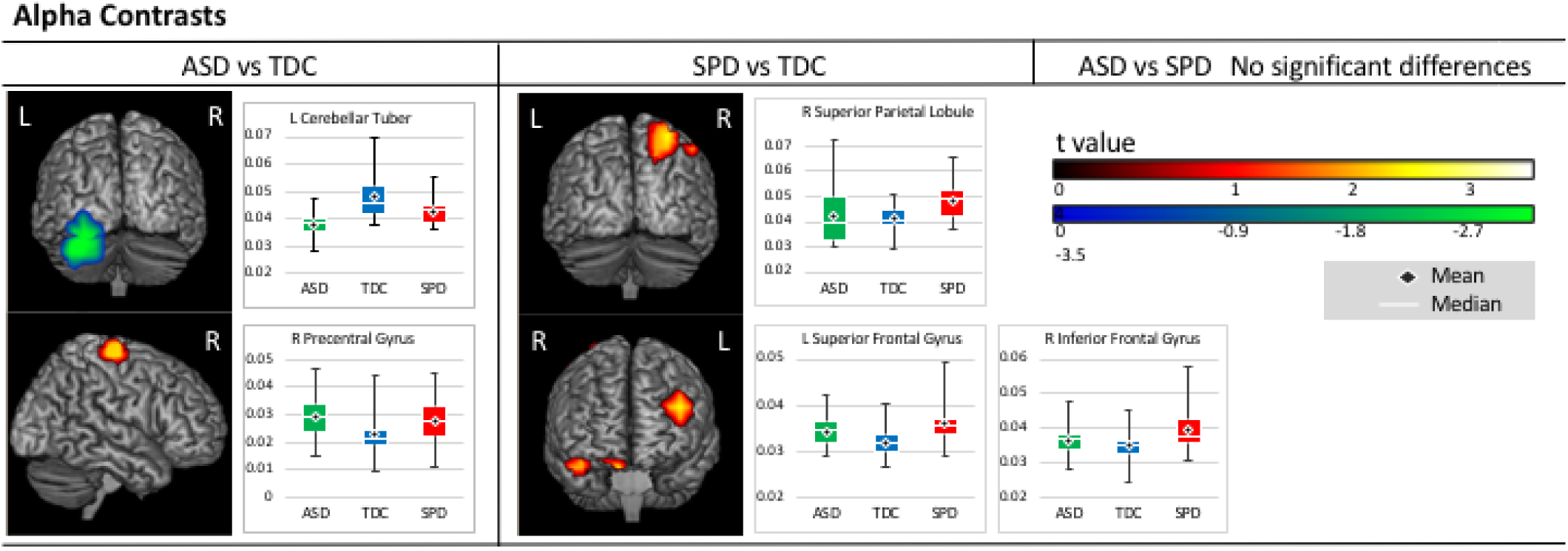
Alpha Contrasts. Areas of significantly increased (warm) and reduced (cool) alpha connectivity are presented on figures for each pairwise contrast. Accompanying boxplots are presented for each cluster showing imaginary coherence values for all groups at the voxel within that cluster that demonstrated the greatest pairwise difference.

**Table 2.**
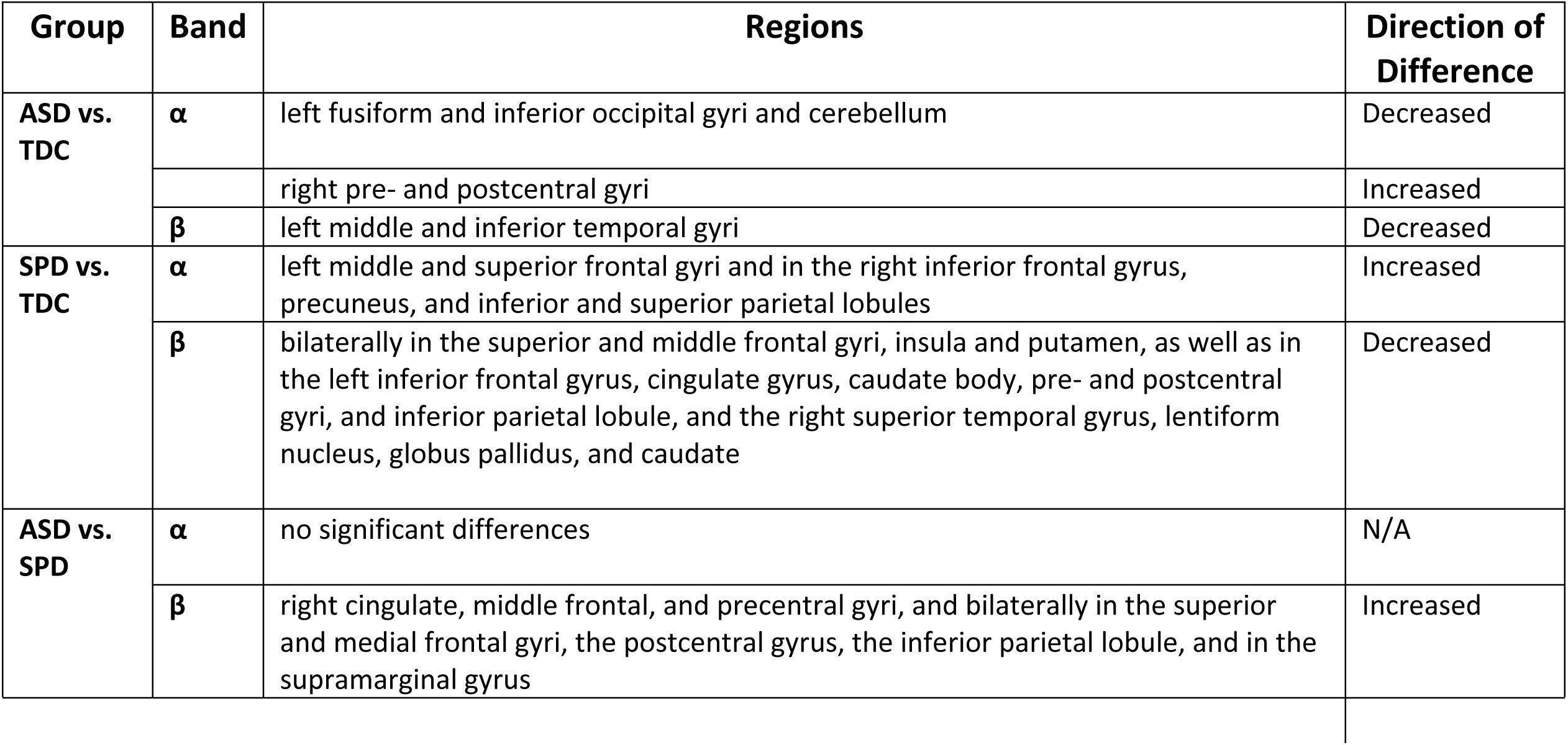
Summary of Group Contrast Results.

### Correlations Between Alpha Connectivity and Sensory Processing/Verbal Abilities

Correlation analyses were performed on all study participants combined across groups to examine the relations between functional connectivity and the range of sensory processing and verbal abilities in our sample. No significant associations were identified between tactile processing performance and measures of alpha coherence; however, significant associations were identified between measures of alpha coherence and auditory processing performance.

Specifically, scores on the DSTP Acoustic scale were positively associated with alpha coherence in the left cerebellar tonsil and negatively associated with alpha coherence in the left inferior and middle temporal gyri. A significant positive association also was identified between VIQ and alpha coherence in the left uncus, cerebellar tonsil, and anterior superior, middle, and inferior temporal gyri (Figure 2). A summary of correlation results is presented in Table 3.

**Figure 2.**
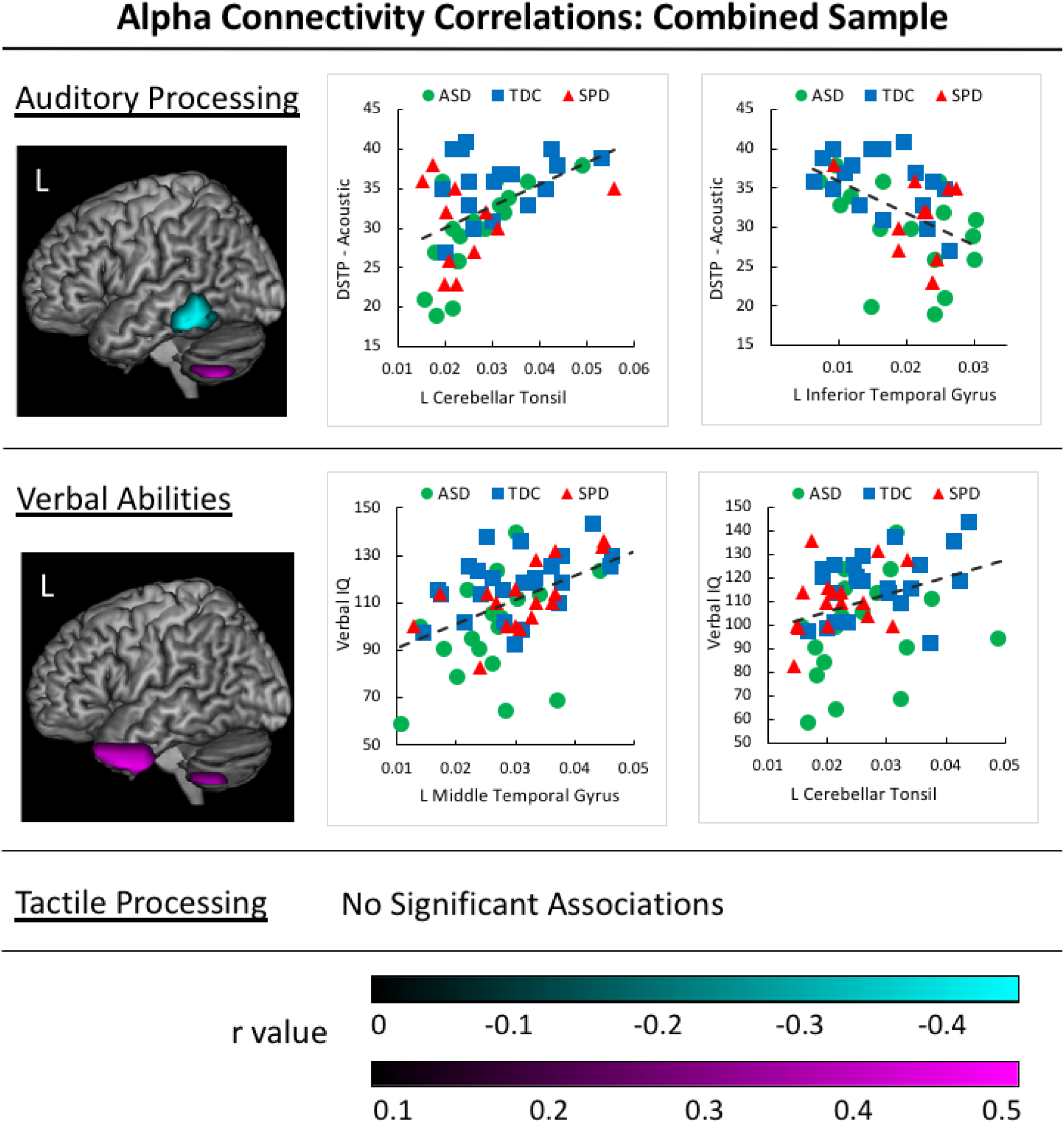
Alpha Correlations in the Combined Participant Sample. Positive associations between auditory processing/verbal abilities and alpha connectivity values are identified in magenta clusters for the sample of all participants in the study. Negative associations between auditory processing and imaginary coherence values are identified in cyan clusters. Corresponding scatterplots are presented for the voxel with the greatest correlation value within each cluster, with groups identified by color and shape (ASD group = yellow circle, SPD group = green triangle, and TDC group = grey square).

**Table 3.** Summary of Correlation Results for the Combine Groups Sample.

| Band | Domain | Task | Regions | Direction of Correlation |
| --- | --- | --- | --- | --- |
| $\alpha$ | tactile | | no significant associations | N/A |
|  | auditory | DSTP Acoustic Scale | left cerebellar tonsil | + |
|  |  |  | left inferior and middle temporal gyri | - |
|  | verbal | VIQ | left uncus, cerebellar tonsil, and anterior superior, middle, and inferior temporal gyri | + |
| $\beta$ | tactile | Graphesthesia | right precentral gyrus | - |
|  | auditory |  | no significant associations | N/A |
|  | verbal |  | no significant associations | N/A |

### Group Contrasts in Beta Connectivity

Group contrasts in beta coherence indicated that, relative to TDC participants, the ASD group showed reduced connectivity in the left middle and inferior temporal gyri. Relative to SPD participants, however, the ASD group showed a pattern of increased beta connectivity in the right cingulate, middle frontal, and precentral gyri, and bilaterally in the superior and medial frontal gyri, the postcentral gyrus, the inferior parietal lobule, and in the supramarginal gyrus. Finally, when compared to TDC participants, the SPD group demonstrated a pattern of reduced beta connectivity bilaterally in the superior and middle frontal gyri, insula and putamen, as well as in the left inferior frontal gyrus, cingulate gyrus, caudate body, pre- and postcentral gyri, and inferior parietal lobule, and in the right superior temporal gyrus, lentiform nucleus, globus pallidus, and caudate. Beta contrasts are presented in Figure 3 and summarized in Table 2.

**Figure 3.**
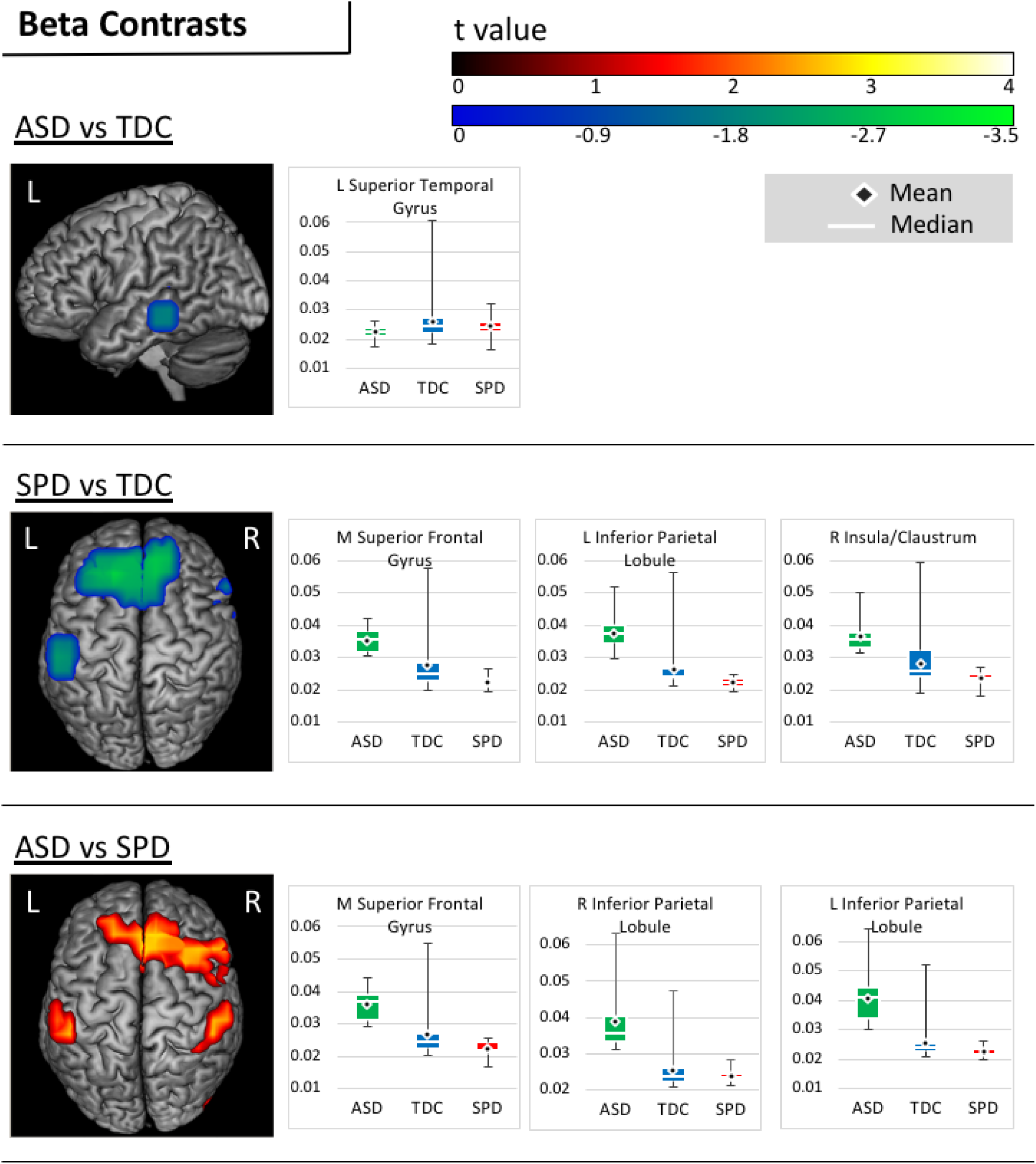
Beta Contrasts. Areas of significantly increased (warm) and reduced (cool) beta connectivity are presented on figures for each pairwise contrast. Accompanying boxplots are presented for each cluster showing imaginary coherence values for all groups at the voxel within that cluster that demonstrated the greatest pairwise difference.

### Correlations Between Beta Connectivity and Sensory Processing

A significant negative association was identified between beta coherence in the right precentral gyrus and performance on the graphesthesia task (Figure 4). No significant associations were identified between beta coherence and measures of auditory processing or verbal abilities in the combined groups sample. A summary of correlation results is presented in Table 3.

**Figure 4.**
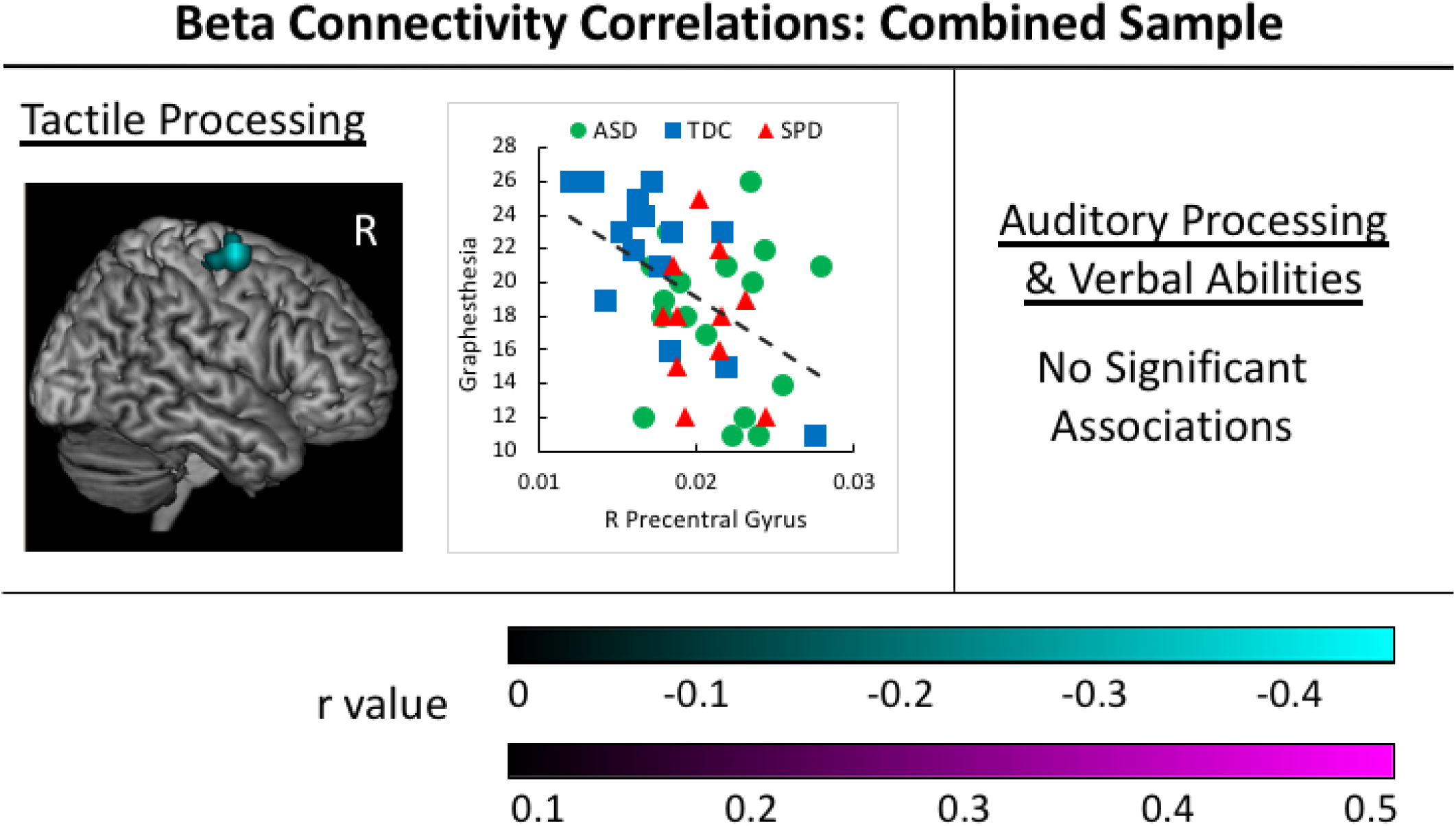
Beta Correlations in the Combined Participant Sample. Negative associations between tactile processing abilities and beta connectivity values are identified in the cyan cluster for the sample of all participants in the study. The corresponding scatterplot is presented for the voxel with the greatest correlation value within each cluster, with groups identified by color and shape (ASD group = yellow circle, SPD group = green triangle, and TDC group = grey square).

## Discussion

This study used two methods to investigate associations between direct assessment of auditory and tactile sensory processing and resting state functional connectivity in the brain. First, we examined differences between groups that would allow us to isolate the sensory processing dysfunction that presents as part of an ASD from that which manifests in the absence of the other defining features of ASD. Second, we directly examined associations between functional connectivity and auditory and tactile processing and verbal abilities in a combined participant sample including all three groups, allowing us to examine the distribution of these variables across children with a range of sensory functioning.

### Group Contrasts in Functional Connectivity

#### ASD vs TDC Contrasts

Relative to the TDC group, participants with ASD showed increased alpha connectivity in the right sensorimotor cortex and decreased connectivity in left posterior fusiform, occipital, and cerebellar regions. Notably, increased alpha power (Edgar, Heiken, et al. 2015) in a similar region in the right medial sensorimotor cortex, and increased alpha to low-gamma phase amplitude coupling in this central midline region (Port et al. 2019) has been reported in prior ASD samples. The present results also recapitulate our previous structural findings in children with ASD, in which we reported decreased structural connectivity in the inferior fronto-occipital fasciculus and the fusiform-hippocampus and fusiform-amygdala tracts (Chang et al., 2014). Our findings of increased cerebellar connectivity are also consistent with considerable prior research implicating the cerebellum in the pathology of ASD. Specifically, cerebellar anomalies, including abnormal anatomy, neurotransmission, oxidative stress, neuroinflammation, and cerebellar motor and cognitive deficits are among the most replicated findings in individuals with ASD (Fatemi et al. 2012).

In the beta range, the ASD group demonstrated decreased beta connectivity in left temporal regions relative to TDC participants. Stronger beta connectivity in TDC relative to ASD participants in temporal regions has been demonstrated in prior work (Kitzbichler et al. 2015). Beta power in the auditory cortex has been hypothesized to be involved in auditory-motor communication (Fujioka et al. 2009) and recent work has demonstrated increases in sensorimotor low beta power in response to perceived self-produced vocal errors on an altered auditory feedback speech paradigm (Franken et al. 2018). The decreased beta connectivity in the left auditory cortex demonstrated in the present study may reflect under-recruitment of this area needed for auditory processing and auditory motor communication in participants with ASD.

#### SPD vs TDC Contrasts

The SPD group differed from the TDC group via increased alpha connectivity in bilateral frontal and right posterior parietal regions and reduced beta connectivity in left parietal and medial and right frontal regions. These differences in functional connectivity identified in these regions may be associated with the impairments in visuomotor skills and attention previously reported in the SPD population (Brandes-Aitken et al., 2018). In fact, prior work examining diffusion imaging in children with SPD identified associations between visuomotor and cognitive control abilities and structural connectivity in regions of the superior longitudinal fasciculus that run adjacent to the parietal regions identified in this study (Brandes- Aitken et al., 2019).

#### ASD vs SPD Alpha and Beta Contrasts

Notably, the ASD and SPD groups did not show significant differences in alpha connectivity. In fact, it was beta connectivity that distinguished these two groups. Specifically, the SPD group showed a pattern of reduced beta connectivity relative to both the TDC and ASD groups in bilateral and medial frontal and left parietal regions. Taken together, these findings suggest that decreased beta connectivity in medial frontal and parietal regions may be involved in, or a response to, the sensory disturbance experienced by children with SPD. Beta activity has been previously reported to be associated with somatosensory gating and attention (Bauer et al. 2012; Buchholz et al. 2014; van Ede et al. 2010). Our previous work has demonstrated common tactile processing deficits in both ASD and SPD groups (Demopoulos, Brandes-Aitken, et al. 2015), although when MEG-acquired somatosensory latencies were compared between these groups, the SPD group demonstrated an intermediate latency and did not significantly differ from TDC or ASD participants (Demopoulos et al. 2017). These previous results, in conjunction with the present finding that beta activity distinguished the ASD and SPD groups in the bilateral somatosensory cortex, may suggest that the pathology underlying tactile dysfunction in these two groups is divergent.

#### Combined Groups Correlation Results

When correlation analyses were performed on all participants combined into one group, alpha connectivity was positively associated with auditory and verbal abilities, whereas beta connectivity was negatively associated with tactile processing. Specifically, there was a common area of positive correlation between left cerebellar alpha connectivity and both auditory processing and verbal abilities; however, an additional positive association was identified between left anterior temporal alpha connectivity and verbal abilities. Previous work has identified an association between increased anterior temporal alpha power and autism symptomatology measured via the SRS total score (Cornew et al. 2012). whereas an additional negative association was identified between posterior temporal alpha connectivity and auditory processing. Taken together, these findings may suggest that increased cerebellar alpha recruitment may be utilized to address auditory processing weakness that affects not only basic auditory processing abilities, a deficit that is common in individuals with ASD (Abdeltawwab and Baz 2015; Alcántara et al. 2004; Demopoulos et al. 2017; Demopoulos, Hopkins, et al. 2015;

Demopoulos and Lewine 2016; DePape et al. 2012; Edgar et al. 2013, 2014; Edgar, Fisk IV, et al. 2015; Gage, Siegel, Callen, et al. 2003; Gage, Siegel, and Roberts 2003; Hitoglou et al. 2010; Järvinen-Pasley and Heaton 2007; Kargas et al. 2015; Oram-Cardy et al. 2005; Oram Cardy et al. 2005; Tecchio et al. 2003; Tomcheck and Dunn 2007), but also verbal abilities. Indeed, prior work has demonstrated links between cortical auditory processing abnormalities and verbal abilities (Berman et al. 2016; Demopoulos et al. 2017; Edgar et al. 2013; Oram-Cardy et al. 2005; Oram Cardy et al. 2008; Roberts et al. 2011, 2012; Schmidt et al. 2009). With regard to beta connectivity, increases in the right somatosensory cortex were associated with poorer performance on the graphesthesia task. Examination of the scatterplot distribution suggests that somatosensory processing limitations may drive the graphesthesia impairments demonstrated in the two clinical groups. Correlation results were consistent with our hypothesis that beta connectivity would be associated with tactile processing and alpha connectivity would be associated with auditory processing. This is consistent with prior work in which alpha oscillations were associated with direction of auditory attention (Bauer et al. 2012) and somatosensory cortex beta band modulation was reported in advance of tactile stimuli (van Ede et al. 2010).

### Limitations and Future Directions

Several limitations of the present study must be acknowledged. First, the participant sample was restricted to males between the ages of 8-12 years. Prior studies examining resting state neural oscillatory behavior have also restricted analyses to males given the high prevalence of ASD in males and sex differences in peak alpha frequency (Edgar et al. 2019; Green et al. 2022; Manyukhina et al. 2022). While these restrictions result in more homogenous groups and minimize confounds of sex and age differences in neurobiology, they also create limitations for the generalizability of these results to females and children and adolescents outside the age range studies. Future research is necessary to understand the applicability of these findings across ages and sexes. This study also included only children with a nonverbal IQ>70, which limits the generalizability of these results to lower functioning individuals. Further, this study focused on only two frequency bands (alpha and beta) and only two sensory domains, auditory and tactile processing. While prior research suggests that these domains may be the most severely impacted in individuals with ASD (Fernandez-Andres et al. 2015), sensory dysfunction is heterogeneous in its presentation among individuals with and without ASD, and understanding neurobiological factors associated with dysfunction in other sensory domains also will be important to inform treatment development. Finally, this study focused on specific aspects of sensory processing (e.g., discrimination, temporal processing, etc.), but did not incorporate measures of sensory responsivity or sensory seeking behavior. Further, this work only focused on two frequency bands, alpha and beta. Future studies could expand upon this work to examine relations between sensory processing dysfunction and functional connectivity in other frequency bands, as gamma oscillatory behavior has been associated with multisensory communication (Misselhorn et al. 2019) and sensory sensitivity (Manyukhina et al. 2021). Future studies are needed to characterize differences in functional connectivity that may account for these heterogeneous sensory responses or behaviors in children with ASD and SPD.

### Conclusions

This study was the first to use MEG to examine participants with ASD and SPD in relation to neurotypical children to identify relevant differences in resting state whole brain functional connectivity that may be associated with sensory dysfunction. This study design allowed us to identify both shared and distinct patterns of neural activity in two groups affected by sensory dysfunction. Specifically, both clinical groups were distinguished from the TDC group by patterns of functional connectivity differences in the alpha and beta bands, whereas the clinical groups were only distinguished from each other on measures of beta connectivity.

Associations between functional connectivity and behavior identified that sensorimotor regions were associated with tactile processing performance and temporal and cerebellar regions were associated with auditory processing and language abilities. These results suggest that resting state differences in oscillatory brain activity in the alpha and beta frequencies is associated with the sensory dysfunction that characterizes children with ASD and SPD.

## Acknowledgments

CD was funded in part by National Institutes of Health grants (K23DC016637- 01A1, R01DC019167-01A1) Autism Speaks CAPD Pilot award 11637, and UCSF Weill Institute for Neurosciences Weill Award for Clinical Neuroscience Research (2016038). EJM was funded by NIH grant K23MH083890, the Wallace Research Foundation and crowdfunding support to the UCSF Sensory Neurodevelopment & Autism Program. SSN was funded in part by National Institutes of Health grants (R01NS100440, R01DC176960, R01DC017091, R01AG062196), UCOP-MRP-17-454755, and the US Department of Defense grant (W81XWH-13-1-0494).

## Author Contribution Statement

C.D., E.J.M., and S.S.N. conceived the project and methodological approach. S.D., S.H., A.F., and D.M. participated in data acquisition. C.D. and X.D. performed data analysis with consultation from A.F., L.B.H. and K.G.R. C.D., X.D., and B.G.J. created the figures. C.D. and M.R.G. created the tables. C.D. wrote the manuscript in consultation with E.J.M. and S.S.N.

## Data Availability Statement

The data that support the findings of this study are available on request from the corresponding author. The data are not publicly available due to privacy or ethical restrictions.

## Additional Information

The authors have no competing interests to declare.

## Notes

### Competing Interest Statement

The authors have declared no competing interest.

